# *Resf1* is required for proper placental development and configuration of trophoblast cell-specific heterochromatin

**DOI:** 10.1101/2025.09.25.678512

**Authors:** Kei Fukuda, Kimiko Inoue, Chikako Shimura, Moe Kitazawa, Michiko Hirose, Shogo Matoba, Atsuo Ogura, Yoichi Shinkai

## Abstract

*Resf1* (Retroelement silencing factor 1) is involved in retroelement silencing cooperated with H3K9 methyltransferase SETDB1 by regulating H3K9 methylation in mouse embryonic stem cells (mESCs). However, it remains unknown whether *Resf1* functions in retroelement silencing *in vivo*, and has a role in development. Here, we established *Resf1*-deficient mice, which exhibited developmental delay, partial embryonic lethality, and placental defects. Notably, retroelements were also upregulated in the *Resf1*-deficient placenta, correlating with increased expression of nearby genes. To further assess whether *Resf1* functions within the trophoblast lineage, we generated *Resf1*-deficient trophoblast stem cell (TSC) lines. Both undifferentiated TSCs and differentiated TSCs (D-TSCs) display increased retroelement expression along with elevated levels of genes associated with placental development. Moreover, *Resf1*-deficient TSCs exhibit compromised maintenance of H3K9me3 domains in a manner independent of SETDB1. Collectively, our findings reveal that *Resf1* plays multifaceted roles beyond retroelement silencing, underscoring its importance in development and its critical function in trophoblast lineage regulation.

## Introduction

Transposons make up approximately half of the mammalian genome, playing a significant role in the acquisition of new genes and the diversification of gene expression regulation, thus contributing significantly to mammalian evolution (Senft and Macfarlan 2021). On the other hand, transposons pose a risk of genome instability, prompting hosts to acquire various defence mechanisms to suppress them (Slotkin and Martienssen 2007; Sekiguchi et al. 2013). Epigenetic mechanisms play a major role in transcriptional repression of transposons, with DNA methylation and H3K9 methylation serving as key components of this silencing process (Matsui et al. 2010; Karimi et al. 2011; Fukuda and Shinkai 2020). The mammalian genome contains a wide variety of transposons, many of which are species-specific (Senft and Macfarlan 2021). To specifically suppress this diverse array of transposons, hosts have evolved various KRAB-ZNF genes that recognize transposon sequences (Imbeault et al. 2017; Senft and Macfarlan 2021). KRAB-ZNF proteins recruit DNA methyltransferases and H3K9 methyltransferases to transposon loci via the adaptor protein TRIM28, thereby repressing their transcription (Rowe et al. 2010; Fukuda and Shinkai 2020).

The mechanisms by which repressive chromatin modifications are induced to and maintained at transposons are complex and vary depending on the type of transposon and its genomic context (Fukuda et al. 2018; Fukuda and Shinkai 2020). Multiple chromatin-associating protein complexes play crucial roles. For example, in mouse embryonic stem (ES) cells, Long Interspersed Nuclear Elements (LINEs) are repressed by the Human Silencing Hub (HUSH) complex, whereas endogenous retroviruses (ERVs) are targeted by the ATRX/DAXX complex (Fukuda et al. 2018; Robbez-Masson et al. 2018; Fukuda and Shinkai 2020). In addition, different H3K9 methyltransferases are involved depending on the genomic regions in which the transposons are inserted (Fukuda et al. 2021; Fukuda et al. 2025a). As described, transposons are mainly repressed by the H3K9 and DNA methylation pathways (Karimi et al. 2011), but when H3K9 methylation is lost, alternative repressive pathways, such as H3K27 methylation and H2A monoubiquitination, can compensate to suppress some transposon activities (Fukuda et al. 2023; Fukuda et al. 2025b).

Previously, we performed CRISPR-Cas9 screening for transposon silencing factors in mESCs and identified numerous novel transposon suppressing factors associating with the SETDB1 pathway (Fukuda et al. 2018). One of the newly identified genes, *Retroelement silencing factor 1 (Resf1)*, which we identified through the screening in mESCs, appears to be primarily involved in H3K9 methylation of ERVs in mESCs (Fukuda et al. 2018). RESF1 interacts with SETDB1 and functions in SETDB1 enrichment on its target (Fukuda et al. 2018). RESF1 also interacts with the core pluripotency proteins OCT4 and NANOG in ESCs (van den Berg et al. 2010; Gagliardi et al. 2013) and supports embryonic stem cell self-renewal (Vojtek and Chambers 2021). *Resf1* has been suggested to play a role in the differentiation of germ cells in *in vitro* differentiation systems (Vojtek and Chambers 2021). However, its roles *in vivo* remain largely unknown, including its potential function in transposon suppression as well as its broader physiological roles in development and cell differentiation. Therefore, we generated *Resf1* knockout (KO) mice and examined above issues.

## Result

### Partial lethality, placental abnormality and developmental delay in *Resf1* KO mice

*Resf1* encodes 1,521 amino acids in mice and is evolutionary conserved at least among mammals. In human, *RESF1* mRNA is highly expressed in the placenta, ovary and spleen, while it is low expressed in the brain (Supplementary Figure S1A). RNA-seq analysis in mouse embryonic day 15.5 (E15.5) placentas and brains exhibit that *Resf1* is also highly expressed in placentas but not in brains in mice (Supplementary Figure S1B). To investigate *in vivo* function of *Resf1*, we generated *Resf1* deficient mice using CRISPR-Cas9 technology. As the third exon of *Resf1* encodes 92% of the protein, we designed two guide RNAs (gRNAs) targeting this exon which can delete most of the exon 3 sequences (Supplementary Figure S1C). By injection of such *Resf1* gRNAs with Cas9 protein into fertilized eggs, we obtained three founders which showed expected size of deletion of exon 3 region of *Resf1*. Since all three founders possessed exactly same deletion at the target locus (Supplementary Figure S1D), we picked one founder for the future analysis. RNA-seq analysis confirmed that truncated *Resf1* mRNA which lost most of the exon 3 was detected in *Resf1* KO placenta (Supplementary Figure S1E). It should be noted, detected *Resf1* mRNA derived from the deleted allele possessed the remained part of exon 3 with the undeleted portion of intron 3, which is normally spliced out. Consequently, exons 4 is connected in-frame. Unfortunately, still no reliable antibody is currently available to detect endogenous RESF1, but such truncated RESF1 may be produced even though 87.5% of the original RESF1 protein sequences are lost.

Then, we examined impact of this *Resf1* KO on mouse development and viability. *Resf1* heterozygous KO (*Resf1*^+/−^) mice showed no clear phenotypes, but when they were intercrossed, the proportion of homozygous KO (*Resf1*^−/−^) pups was less than 5% in both natural mating and in vitro fertilization (IVF), which is markedly lower than the expected Mendelian ratio (Figure 1A, B). Surviving *Resf1*^−/−^ mice exhibited a body weight approximately 20–30% lower than that of other genotypes, and this reduction persisted into sexual maturity (Supplementary Figure S2A, B). Loss of *Resf1* in mouse embryonic stem cells (mESCs) has been reported to impair self-renewal and reduce the efficiency of germline entry (Vojtek and Chambers 2021). To explore whether *Resf1* deficiency also affects fertility *in vivo*, we assessed the reproductive performance of *Resf1*^−/−^ mice by mating them with *Resf1*^+/−^ partners. While *Resf1*^−/−^ males were capable of producing offspring at normal rates, *Resf1*^−/−^ females exhibited reduced fertility, as evidenced by lower birth rates and smaller litter sizes compared to other genotype pairings (Supplementary Figure S2C, D). However, histological analysis of the ovaries revealed the presence of normally matured follicles in *Resf1*^−/−^ females (Supplementary Figure S2E). Moreover, successful fertilization was observed through IVF using *Resf1^−/−^*oocytes, and they could implant normally (Supplementary Figure S2F). These findings suggest that while *Resf1*^−/−^ oocytes are capable of being fertilized and undergoing implantation, the overall success rate of live birth is markedly reduced. Therefore, the absence of RESF1 impairs female reproductive performance, particularly at the post-implantation stage.

**Figure 1.**
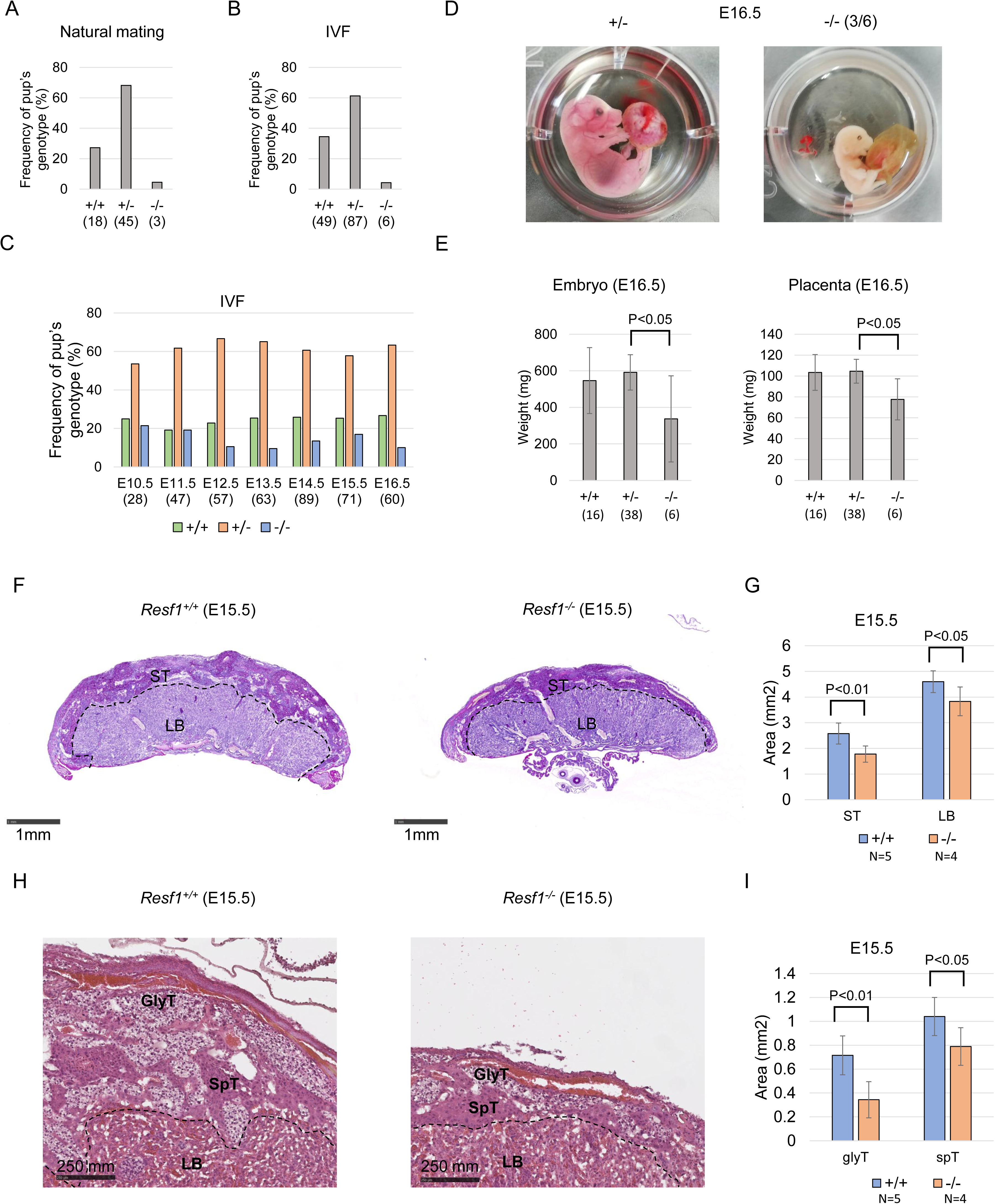
Partial embryonic lethality of *Resf1* KO mice. (A, B) Genotyping of offspring resulting from the cross between *Resf1* heterozygous mice. Both in natural mating and in vitro fertilization (IVF), the rate of homozygous KO pups was lower than predicted by Mendelian ratios. The number of pups investigated is indicated in parentheses. (C) Genotyping of the fetuses resulting from the cross between *Resf1* heterozygous mice in each embryonic day. The decreased rate of *Resf1* KO fetuses begins from E12.5. The number of fetuses investigated is indicated in parentheses. (D) Representative image of E16.5 *Resf1* KO fetuses. Half of E16.5 *Resf1* KO fetuses are pale and small. (E) Weight of fetus and placentas at E16.5. *Resf1* KO fetuses exhibit lighter weights for both fetuses and placentas compared to other genotypes. The P-values were calculated using a Student’s T-test. (F) PAS staining of E15.5 WT and *Resf1* KO placentas. Scale bar indicates 1mm of length. ST and LB denote spongiotrophoblast layer and labyrinth layer, respectively. (G) Area of ST and LB in WT or *Resf1*^−/−^ E15.5 placentas. Both ST and LB are smaller in *Resf1*^−/−^ placentas compared to WT, with the ST being more severely affected. The P-values were calculated by Student’s T-test. (H) HE stains of E15.5 WT and *Resf1*^−/−^ ST. GlyT and SpT correspond to glycogen trophoblast and spongiotrophoblast, respectively. The dashed line indicates the boundary between LB and ST. Scale bar indicates 250 mm of length. (I) Area of GlyT and SpT in WT or *Resf1*^−/−^ E15.5 placentas. Both area ST and LB are smaller in *Resf1*^−/−^ than WT, but glyT is more severely affected. The P-values were calculated by Student’s T-test.

Next, we addressed the potential cause of the low frequency of homozygous offspring resulting from intercrosses between *Resf1* heterozygous mice. The decrease in the ratio of *Resf1^−/−^*individuals was observed from embryonic day 12.5 (E12.5) onward (Figure 1C). Among the fetuses observed at E16.5, half of them were undersized and exhibited a pale phenotype (Figure 1D, E). Additionally, not only was the fetuses itself affected, but the placental weight was also reduced in the *Resf1* KO fetuses (Figure 1E). Histological analysis of E15.5 placenta revealed significant smaller Spongiotrophoblast layer in *Resf1*^−/−^ placenta than WT samples (Figure 1F, G). Furthermore, there was a significant reduction in the amount of Glycogen trophoblasts within the Spongiotrophoblast layer in the *Resf1^−/−^* placentas as compared to WT samples (Figure 1H, I). Reduction of Spongiotrophoblast layer and Glycogen trophoblasts were also observed in E13.5 *Resf1^−/−^* placenta (Supplementary Figure S3A). In the mouse placenta, multinucleated syncytiotrophoblast cells form via the fusion of trophoblast cells between E10.5 and E16.5. This process leads to a thinner barrier between maternal and fetal blood, thereby increasing the surface area available for maternal-fetal exchange of gases, nutrients, and waste products. This phenomenon holds significant importance in placental morphogenesis (Huppertz et al. 2001; Georgiades et al. 2002; Bischof and Irminger-Finger 2005; Watson and Cross 2005; Dupressoir et al. 2009). However, in *Resf1^−/−^*mice, the placentas exhibited clusters of densely packed trophoblast cells, and the spaces within fetal blood vessels were significantly reduced (Supplementary Figure S3B). These observed clusters are thought to arise due to the incapability of trophoblast cells to undergo fusion or decreased placental vascularization. These results suggest that intact *Resf1* is required for normal placental development.

### Dysregulation of endogenous retrovirus in *Resf1^−/−^* placenta

To investigate the underlying causes of the placental malformations observed in *Resf1^−/−^* mice, we performed RNA-seq analysis on E15.5 placentas to identify gene expression abnormalities. 60 genes and 24 genes showed significantly higher and lower in *Resf1^−/−^*placentas compared to WT, respectively (Figure 2A). Among the upregulated genes, Gene Ontology terms related to cytolysis (FDR = 3.0 × 10⁻³) and peptidase activity (FDR = 1.3 × 10⁻²) were significantly enriched. The enrichment of Peptidase activity is largely due to granzyme gene cluster (*Gzmc, Gzmd*). Other gene clusters, including Prolactin gene clusters, were also upregulated in *Resf1^−/−^* placentas (Figure 2A).

**Figure 2.**
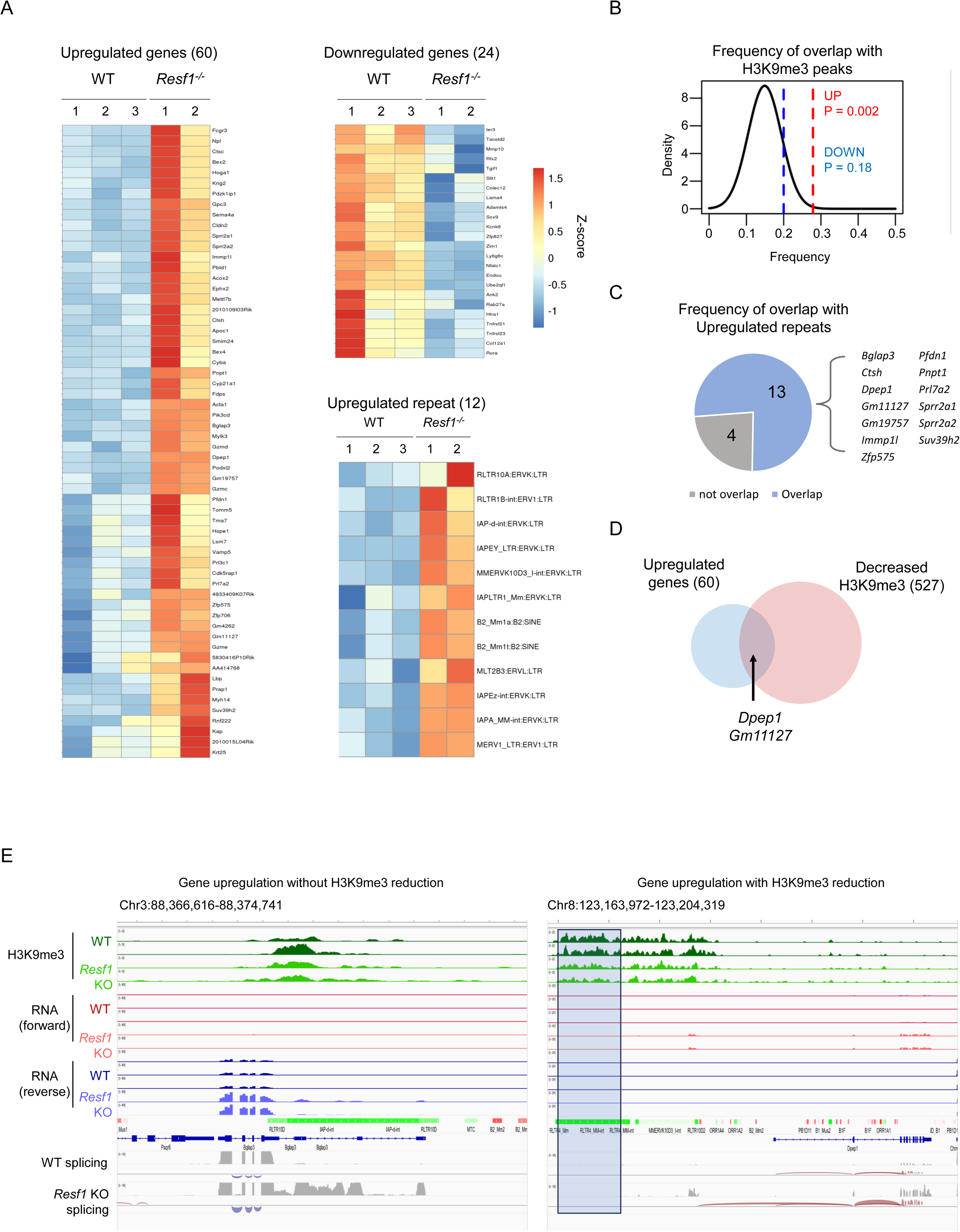
Transcriptome analysis of E15.5 WT and *Resf1*^−/−^ placentas. (A) Heatmap of Z-scaled expression levels of differentially expressed genes and repeats. LTR-type transposons are mainly upregulated in *Resf1*^−/−^ placentas. (B) Frequency of differentially expressed genes overlapping with H3K9me3 peaks. The frequency indicates the presence of H3K9me3 peaks in wild-type samples within 20 kb upstream and 5 kb downstream of the transcription start site. The p-value was calculated using a random permutation test. (C) Frequency of H3K9me3 peaks near upregulated genes that overlap with upregulated repeats. (D) Overlap between differentially expressed genes and decreased H3K9me3 peaks. (E) Representative examples of transposon-derived transcription associated with expression upregulation: one with decreased H3K9me3 and the other without.

We also identified 12 upregulated transposons in the *Resf1^−/−^*placenta and the majority of them is LTR type transposons, which is consistent with our previous results in *Resf1^−/−^* mESCs (Fukuda et al. 2018). H3K9me3 ChIP-seq analysis in WT E15.5 placenta showed that upregulated genes were significantly enriched around H3K9me3 peaks (17/60, P=0.002) (Figure 2B) and H3K9me3 peaks around upregulated genes were frequently overlapped with upregulated repeats (13/17) (Figure 2C). To investigate whether the gene expression changes observed in the placenta are associated with alterations in H3K9me3, we performed ChIP-seq analysis for H3K9me3 in E15.5 placentas from both WT and *Resf1*^−/−^ mice. As a result, we identified 736 regions with increased H3K9me3 and 527 regions with decreased H3K9me3 in *Resf1^−/−^*placentas (Supplementary Figure S4A). Consistent with the role of *Resf1* in suppressing LTR-type transposons, the regions with reduced H3K9me3 were enriched for LTR transposons (Increased regions: 41.8%, Decreased regions: 70.2%) (Supplementary Figure S4B). However, regions with reduced H3K9me3 were identified only around two upregulated genes (Figure 2D). Even if transcription initiated from transposons contributes to the upregulation of these genes, the presence or absence of changes in H3K9me3 varied among them (Figure 2E). Therefore, in the placenta, the suppression of genes and transposons by *Resf1* may not necessarily be due to the direct regulation of H3K9me3 near their TSS regions.

### Endogenous retrovirus dysregulation in *Resf1^−/−^* TSCs and D-TSCs

Considering the diverse cell types present in the placenta, including those from the hematopoietic origin and maternal tissues, we established three independent female Trophoblast Stem Cells (TSC) lines each from *Resf1*^+/−^ and *Resf1*^−/−^ embryos, in order to investigate potential abnormalities intrinsic to the trophoblast lineage (Figure 3A). We differentiated all established TSC lines (D-TSC) and investigated the effect of *Resf1* loss in transcriptome and H3K9me3 using RNA-seq and H3K9me3 ChIP-seq (Fig. 3B, Supplementary Figure S5A). We also established two *Resf1^+/−^* and one *Resf1^−/−^* ESC lines (Figure 3A). RNA-seq analysis of TSCs, D-TSCs, and ESCs revealed unique gene expression profiles that clustered according to cell type, with *Resf1*^−/−^ and *Resf1*^+/−^ samples also exhibiting distinct patterns from each other (Figure 3C). Additionally, examining the expression of marker genes for each cell type revealed that ESCs, TSCs, and D-TSCs expressed their respective marker genes (Figure 3D). Notably, D-TSCs expressed marker genes for Glycogen Trophoblast (GlyT), Syncytiotrophoblast (SynT), and Trophoblast giant cell (TGC), indicating differentiation into various cell types (Figure 3D).

**Figure 3.**
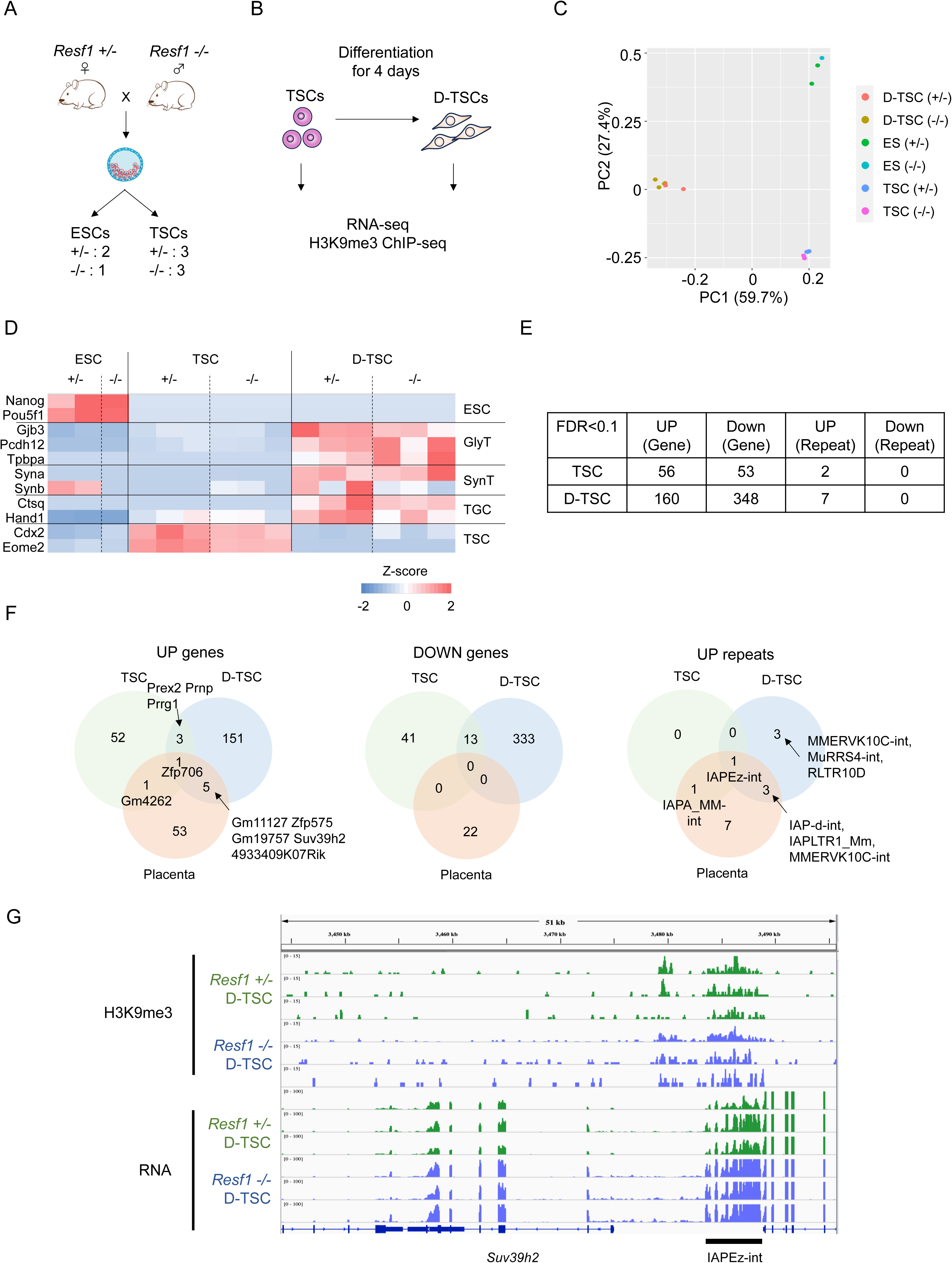
Transcriptome analysis of TSCs and D-TSCs. (A) Schema of establishment of *Resf1*^−/−^ ESCs and TSCs. (B) Experimental schema of TSC differentiation and NGS analysis. (C) PCA analysis of gene expression profiles among samples. (D) Marker gene expression profiles in each cell line. (E) Number of differentially expressed genes and repeats in *Resf1*^−/−^ TSCs and D-TSCs. (F) Venn diagrams of differentially expressed genes and repeats among TSCs, D-TSCs and placentas. (G) Representative region of differentially expressed gene located near retroelement.

We identified 109 (UP: 56, DOWN: 53) and 508 (UP: 160, DOWN: 348) differentially expressed genes in TSCs and D-TSCs, respectively (Figure 3E). In addition, two repeats and seven repeats showed increased expression in *Resf1*^+/−^ TSCs and D-TSCs, respectively, and all of them belonged to the LTR-type transposons (Figure 3E, Supplementary Table S1). Most of all dysregulated genes in TSCs, D-TSCs and placentas had not overlapped each other (Figure 3F). For transposons, only IAP was commonly upregulated among TSC, D-TSCs and placentas (Figure 3F). Thus, it became evident that the genes and transposons regulated by *Resf1* vary significantly across tissues and cell types. Upstream of the *Suv39h2* gene lies an IAPEz-int element. In *Resf1* KO D-TSCs, transcription is initiated from this IAPEz-int, leading to increased expression of *Suv39h2*. Although H3K9me3 is normally enriched at this IAPEz-int, no reduction is observed upon *Resf1* KO (Figure 3G). Similar to the placenta, this finding indicates that in the trophoblast lineage, a decrease in H3K9me3 is not necessarily required for transcriptional activation in *Resf1* KO cells.

The differentially expressed genes were enriched with various GO terms, including those associated with development, differentiation, and notably, terms related to angiogenesis (Supplementary Figure S5B). Interestingly, this accumulation suggests a potential connection to blood vessel formation. Given the suspected developmental defects in fetal blood vessels in *Resf1*^−/−^ placenta, there might be a correlation between these findings.

### Instability of H3K9me3 domain maintenance in *Resf1^−/−^* TSCs

To investigate whether *Resf1* regulates H3K9me3 also in the trophectoderm lineage, we first identified differentially enriched peaks (DE peaks) between WT and KO samples. In TSCs, we found 1,423 regions with decreased H3K9me3 levels and 458 regions with increased levels upon *Resf1* KO. In D-TSCs, 1,426 decreased and 717 increased regions were identified (Supplementary Figure S6A). Then, we compared DE peaks between TSCs and D-TSCs and found 481 decreased peaks and 58 increased peaks were shared, indicating a higher degree of overlap among the decreased peaks. In regions where H3K9me3 changes occur in TSCs, H3K9me3 levels are comparable between TSCs and D-TSCs. In contrast, in regions where H3K9me3 changes occur in D-TSCs, mild alterations are already detectable in TSCs, but these changes become more pronounced upon differentiation (Supplementary Figure S6B). Taken together, these results indicate that H3K9me3 alterations are more prominent in D-TSCs than in TSCs.

Previously we reported that *Resf1* is involved in SETDB1-mediated retroelement silencing (Fukuda et al. 2018). To investigate whether the reduction of H3K9me3 in *Resf1^−/−^* TSCs is due to defect of the SETDB1-mediated H3K9me3 pathway, we generated TSCs harboring a doxycycline-inducible shRNA targeting *Setdb1* or *LacZ* mRNAs. Having successfully knocked down *Setdb1* in these cell lines, we performed H3K9me3 ChIP-seq analysis using these cells (Supplementary Figure S7). From these ChIP-seq data, we identified 3,994 decreased DE peaks and 428 increased DE peaks in *Setdb1* knockdown (KD) TSCs (Supplementary Figure S7B). Only five regions overlapped between the decreased DE peaks in *Resf1*^−/−^ TSCs and those in *Setdb1* KD TSCs (Supplementary Figure S7C). Moreover, whereas more than 10% of the decreased DE peaks in *Setdb1* KD were located within IAP elements, fewer than 2% of such peaks in *Resf1*^−/−^ TSCs or D-TSCs showed this association. (Supplementary Figure S7D). In addition, the regions surrounding the decreased DE peaks in *Setdb1* KD TSCs exhibited globally low levels of H3K9me3, whereas those in *Resf1*^−/−^ TSCs/D-TSCs showed high H3K9me3 levels in the surrounding regions that were reduced upon *Resf1* KO (Supplementary Figure S6C and S7B). These results indicate that even though *Resf1* was identified as a gene associating with SETDB1-mediated transposon silencing function, the H3K9me3 regions regulated by *Setdb1* and those regulated by *Resf1* have distinct characteristics, suggesting that in TSCs, *Resf1* regulates H3K9me3 through a *Setdb1*-independent pathway.

Consistent with the observation that regions surrounding the DE peaks in TSCs also showed a reduction in H3K9me3, the decreased DE peaks tended to form clusters, and H3K9me3 reduction was observed across large domains spanning megabase (Mb)-scale regions (Supplementary Figure S6D). In TSCs, it has been reported that Mb-scale H3K9me3 domains are formed and can act as a barrier to reprogramming during somatic cell nuclear transfer (Hada et al. 2022). These findings suggest that *Resf1* may be involved in the formation of such H3K9me3 domains. To capture broader changes in H3K9me3 beyond narrow peaks, we identified H3K9me3 domains using *Hiddendomains*, subdivided each domain into 100-kb bins, and detected bins showing changes in H3K9me3 levels (Starmer and Magnuson 2016). In TSCs and D-TSCs, *Resf1* KO resulted in a reduction of H3K9me3 in 101 and 1,630 domains, respectively (Figure 4A, Supplementary Figure S8A). Only one upregulated gene was located near decreased H3K9me3 domains (Supplementary Figure S8A). Therefore, it can be inferred that the decrease in H3K9me3 due to *Resf1* deficiency is largely unrelated to the upregulation of gene expression. Unlike decreased H3K9me3 domains, only 3 and 98 H3K9me3 domains showed increased H3K9me3 enrichment in TSCs and D-TSCs (Supplementary Figure S8A). As most of H3K9me3 decreased domains in TSCs (97/101) were also showed differential H3K9me3 enrichment in D-TSCs (Supplementary Figure S8A), the following text primarily focuses on decreased H3K9me3 domains in D-TSCs.

**Figure 4.**
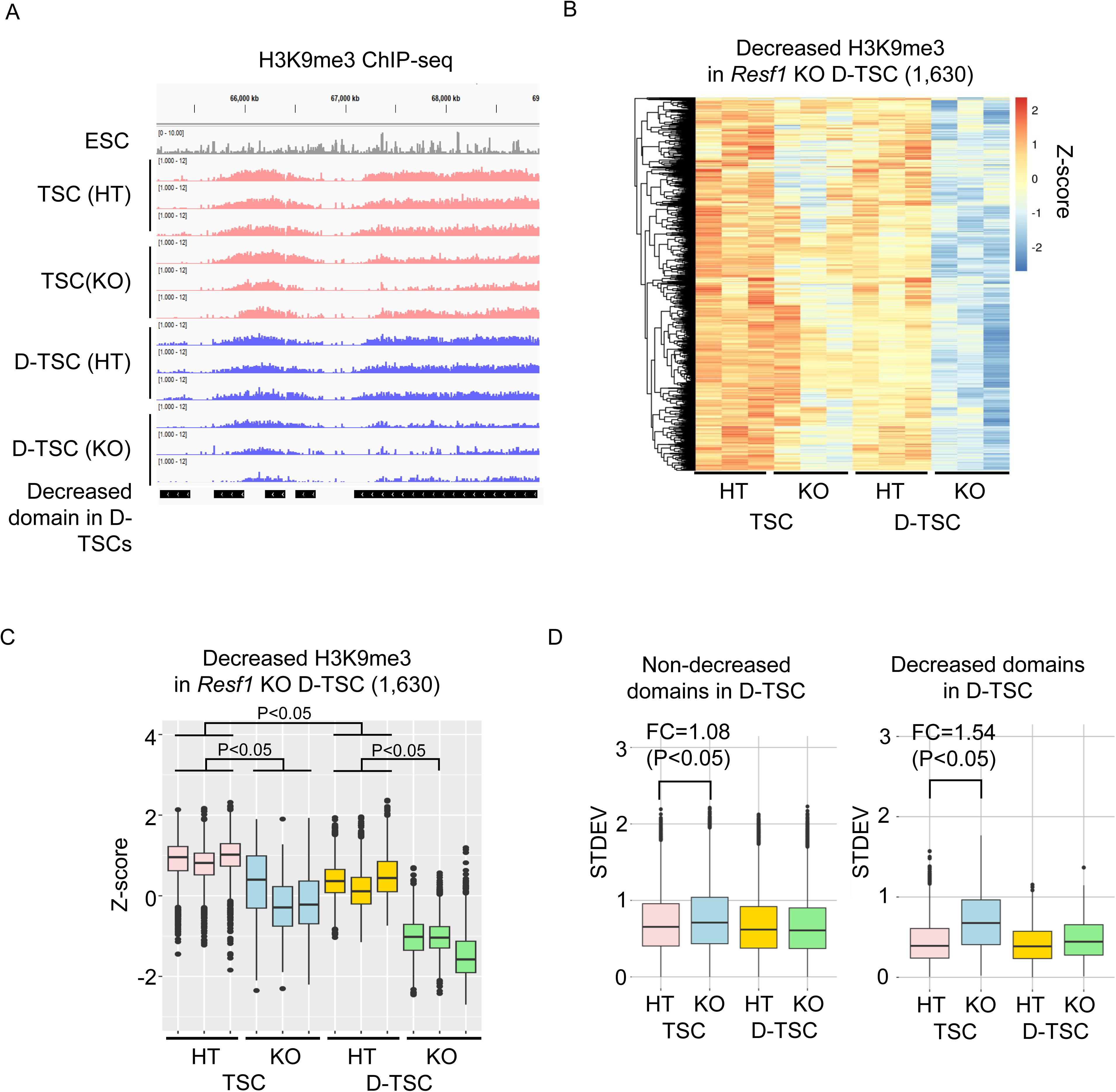
Destabilization of H3K9me3 domains in *Resf1* KO TSCs. (A) Representative view of decreased H3K9me3 domains in D-TSCs. Black boxes represent decreased H3K9me3 domains. HT and KO denote *Resf1^+/−^* and *Resf1^−/−^*, respectively. (B) Heatmap of H3K9me3 levels in decreased H3K9me3 domains. Heatmap is colored by Z-scored H3K9me3 enrichment. (C) Boxplots of H3K9me3 enrichment in decreased H3K9me3 domains. The regions where H3K9me3 decreases in *Resf1*^−/−^ D-TSCs exhibit a tendency of reduced levels in TSCs as well, and furthermore, these are regions that show decreased levels during TSC differentiation. P-values are calculated by Tukey’s test. (D) Variation of H3K9me3 enrichments among clones. Boxplots showed STDEVs of H3K9me3 enrichment among clones in non-DE regions (left) and decreased regions (right) in D-TSCs. *Resf1* KO TSCs exhibit significant variation among clones in the decreased regions. P-values are calculated by Student’s T-test.

**Figure 5.**
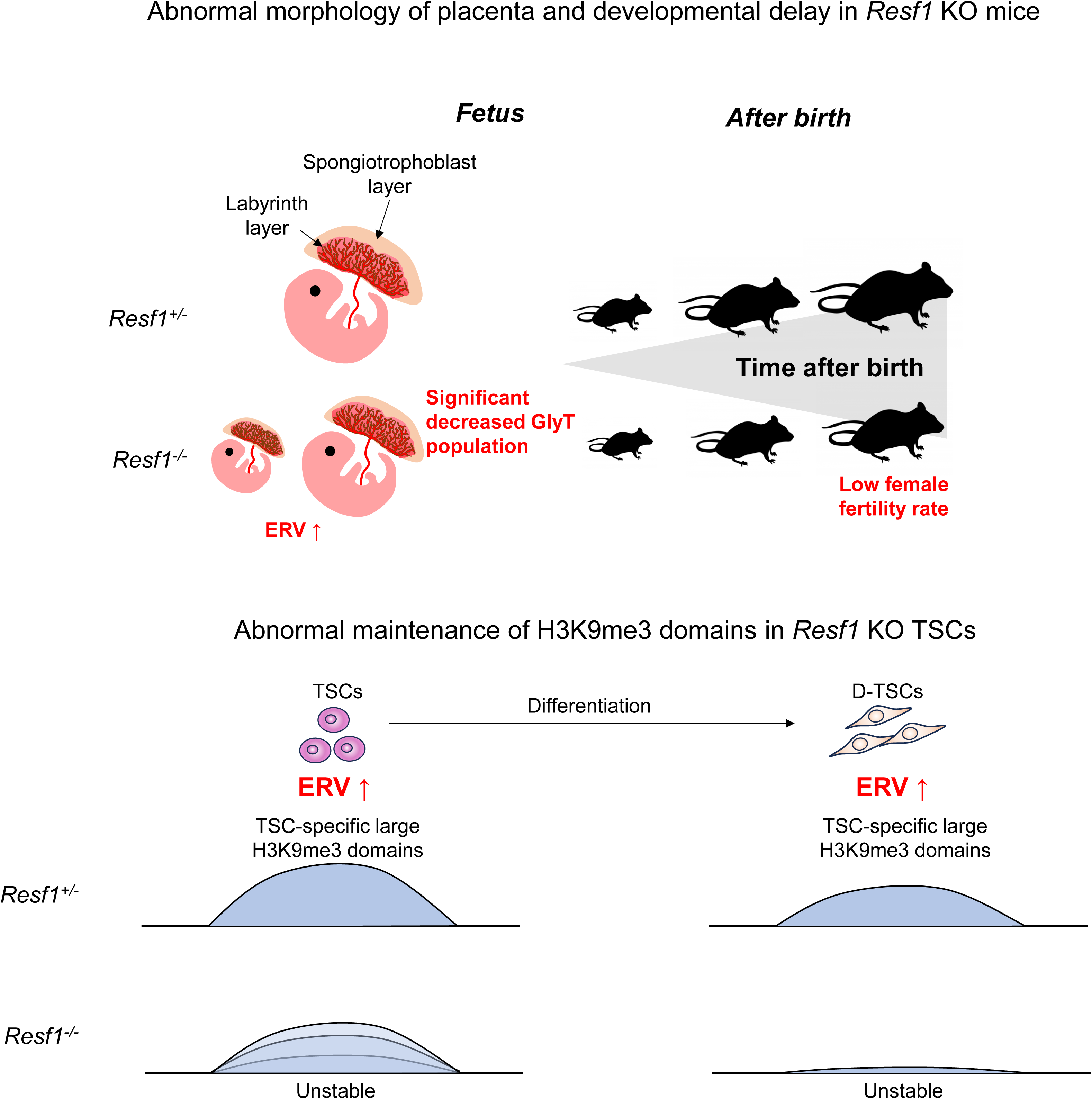
Summary of *Resf1* KO phenotype. *Resf1* KO mice exhibit partial lethality accompanied by placental insufficiency, and postnatally, they continue to display growth delay. Furthermore, within the trophoblast lineage, there is an increase in endogenous retrovirus expression and destabilization of heterochromatin.

Length distribution of decreased/increased/non-different H3K9me3 domains showed most of decreased H3K9me3 domains are 100-kb which is the maximum length of H3K9me3 domains (Supplementary Figure S8B). Contrary to upregulation of LTR-type transposons by *Resf1* deficiency, decreased H3K9me3 domains were AT-rich/LINE-rich and SINE/LTR-poor (Supplementary Figure S8C, D). Comparison of H3K9me3 enrichment in decreased H3K9me3 domains in *Resf1^−/−^* D-TSCs among samples revealed that H3K9me3 decreases during differentiation in *Resf1*^+/−^ cells. *Resf1*^+/−^ TSCs had higher H3K9me3 levels than *Resf1*^+/−^ D-TSCs (Figure 4B, C). In *Resf1*^−/−^ TSCs, H3K9me3 levels were lower than in *Resf1*^+/−^ TSCs, but both the extent of reduction and the affected regions were heterogeneous among clones. (Figure 4B–D). These results suggest that *Resf1* plays a particularly important role in maintaining H3K9me3 in regions where its stability weakens during TSC differentiation.

Large H3K9me3 domains in TSCs are formed on H3.1-rich regions mediated by *Chaf1* (Hada et al. 2022). To investigate whether the loss of *Resf1* also affects H3.1 levels, we performed H3.1 ChIP-seq in *Resf1*^+/−^ and *Resf1*^−/−^ TSCs. However, we found that H3.1 levels were not necessarily reduced in the H3K9me3-decreased domains (Supplementary Figure S9A, B). These results indicate that the reduction of H3K9me3 upon *Resf1* deficiency occurs independently of H3.1 loss. Therefore, *Resf1* likely acts downstream of H3.1 deposition.

In this study, we demonstrated that *Resf1* plays a critical role in suppressing LTR-type transposons in the placenta and trophectoderm lineage and is essential for proper placental development. Furthermore, we uncovered a novel function of *Resf1* in regulating large H3K9me3 domains through a *Setdb1*-independent pathway.

## Discussion

In this study, we generated *Resf1* KO mice to investigate the *in vivo* function of a retroelement-suppressing factor *Resf1* and examined their phenotypes. The KO resulted in partial embryonic lethality and growth retardation (Figure 1A, B, Supplementary Figure S2A, B). In addition, KO females exhibited partial infertility (Supplementary Figure S2C, D). Since normal ovarian follicles were observed, oocyte development itself may not be severely defective (Supplementary Figure S2E). However, it has been reported that KO of *Resf1 in vitro* reduces the efficiency of differentiation from embryonic stem cells to germ cells (Vojtek and Chambers 2021), raising the possibility that oocyte development efficiency may be reduced. In addition to partial embryonic lethality, loss of *Resf1* resulted in growth retardation and structural abnormalities of the placenta (Figure 1, Supplementary Figure S3). In mESCs, *Resf1* is involved in the regulation of imprinted genes (Fukuda et al. 2018). Consistent with this, in our present study, we observed upregulation of *Igf2* in TSCs, downregulation of *H19* in D-TSCs, and upregulation of *Peg10* in D-TSCs (Supplementary Table S1). Because imprinting defects are often associated with placental abnormalities (Coan et al. 2005), these aberrant expression patterns of multiple imprinted genes may underlie the placental defects observed in *Resf1*-deficient mice. In addition, deficiency of another imprinted gene, *Rtl1*, leads to partial embryonic lethality due to placental malformation and impaired maintenance of placental structure (Sekita et al. 2008; Kitazawa et al. 2017). *Rtl1* expression was reduced in one *Resf1^−/−^* placenta: all samples showed log2(TPM) values of 4.5 or higher, except for one KO individual, which had a value of 3.3, suggesting inter-individual variability in imprinting regulation in *Resf1*^−/−^ mice. A similar increase in variability was also observed in the H3K9me3 state in TSCs, with greater variation in KO samples than in heterozygous samples (Figure 4D). Such variability may contribute to inter-individual differences in phenotypic outcomes, such as partial embryonic lethality.

Loss of *Resf1* led to increased expression of LTR-type retrotransposons across placenta, TSCs, and D-TSCs. In particular, IAPEz-int showed elevated expression in all three cell types (Figure 3F). Although H3K9me3 is enriched in transposons exhibiting increased expression in *Resf1^−/−^*, the upregulation of expression did not always correlate with a decreased in H3K9me3 levels. Several possibilities may explain this observation. One is that *Resf1* functions downstream of H3K9me3 to repress transcription. Another is that short-read ChIP-seq has limited ability to quantify the expression and H3K9me3 status of individual transposon copies. If *Resf1* deficiency causes upregulation and H3K9me3 loss only in a subset of copies, detecting such changes is challenging. This issue might be addressed by applying long-read sequencing-based RNA and ChIP analyses. Although the underlying mechanism remains unclear, it is a fact that transposon expression increases upon *Resf1* deficiency. Whether this upregulation is directly linked to placental developmental defects is still unknown. However, increased transposon expression often triggers immune responses and induces inflammation, which could potentially contribute to abnormal placental development or dysfunction.

*Resf1* was originally identified as a factor involved in the SETDB1-dependent H3K9me3 pathway. In this study, we newly demonstrated that *Resf1* also contributes to H3K9me3 domains independent of *Setdb1*. In mice, H3K9me3 is mediated not only by *Setdb1* but also by *Suv39h1* and *Suv39h2*, suggesting that *Resf1* might be involved in the *Suv39h1/2* pathway. Indeed, *Suv39h1/2* forms H3K9me3 domains in AT-rich, LINE-rich regions that correspond well with *Resf1*-dependent H3K9me3 domains. However, whether *Resf1* directly participates in this methylation remains unclear and requires further investigation. H3K9me3 domains observed in TSCs are absent in ES cells and E15.5 placenta but are present in extraembryonic tissues (Wang et al. 2018). In humans, the presence of H3K9me3 domains has also been reported in second-trimester cytotrophoblasts (Zhang et al. 2021). The failure to identify such domains in E15.5 mouse placenta in our current study may be due to the heterogeneous mixture of various cell types, including blood cells. By isolating and analyzing specific cell populations, it may be possible to determine which cell types rely on *Resf1* for H3K9me3 domain formation.

This study revealed a novel role of *Resf1* in chromatin regulation and suggested its function *in vivo*, as well as its potential contribution to phenotypic robustness. Thus, *Resf1* may play an important role in maintaining phenotypic stability. Dysregulation of this control system could increase phenotypic variability, potentially elevating the risk of diseases and infertility. Therefore, further studies are needed to elucidate the molecular mechanisms by which *Resf1* ensures phenotypic robustness and to explore its implications for disease pathogenesis and reproductive health. However, although the *Resf1* KO mice and cells used in this study lack almost the entire coding region, some truncated protein products might remain. These residual proteins could exert dominant-negative effects, contributing to the observed phenotypes. This possibility requires further validation in future investigations.

## Materials and Methods

### Animals

Mice used in this study were under C57BL6/J background and housed as a group (4/cage) in 12 hours light/dark cycle. All mice experiments were approved by RIKEN safety divisions and conducted under institute rules and regulations.

### Establishment of *Resf1*-deficient mice

To delete exon 3 of *Resf1*, two crRNAs (UAUCUCAUAAUCCCGACGGAguuuuagagcuaugcuguuuug, AAAACCAAUAGUUGGGCCCGguuuuagagcuaugcuguuuug, 30.5 pmol/ml each), tracrRNA (30.5 pmol/ml) (FASMAC) and Cas9 (M0646, NEB) were injected into nucleus of fertilized eggs. A total of 172 embryos were injected and transplanted, yielding 32 offspring, among which three males carried a large deletion in the *Resf1* exon 3. Since the deletion occurred at the same position in all three males, one individual was selected for subsequent analysis. The following primers were used for genotyping PCR: TGGCCTCACAGTCTTCAATG, GATTAAGTCCCTGACTTCCTTGATGTG, GTCTAATACCAGTGGTTTGACCACCA.

### Establishment of *Resf1*-deficient ESCs and TSCs

ESC and TSC lines were established from blastocysts generated by IVF using oocytes from *Resf1+/-* female mice and spermatozoa from *Resf1*−/− male mice. For establishment of ESCs, developing IVF-derived blastocysts were treated with acidic Tyrode’s solution (Merk Millipore) to remove the zona pellucida, and then transferred directly onto feeder cells in ESC medium containing knock-out DMEM (Gibco), 15% knockout serum replacement (KSR; Thermo Fisher), MEM nonessential amino acid solution (Gibco), GlutaMAX supplement (Gibco), 100 μM 2-mercaptoethanol (2-ME; Merk Millipore), ESGRO recombinant mouse LIF protein (Merk Millipore), 1 μM PD0325901 (Stemgent), and 3 μM CHIR99021 (Stemgent). After several passages, cell lines with typical ESC colonies were selected and used for further analysis.

For establishment of TSCs, developing blastocysts were transferred directly onto feeder cells in chemically defined medium (CDM). CDM was prepared with Neurobasal medium (Gibco, 21103-049) and DMEM high glucose (Sigma, D6429), supplemented with N2 supplement (1×; Gibco, 17502-048), B27 supplement (1×; Gibco, 12587-610), penicillin– streptomycin (1×), 2-mercaptoethanol (150 µM; e-Nacalai), bovine serum albumin (0.05%), KnockOut Serum Replacement (1%; Gibco), recombinant mouse basic fibroblast growth factor (50 ng/ml; Wako, 450-33), recombinant human activin A (20 ng/ml; R&D Systems, 338-AC), XAV939 (10 µM; Calbiochem), Y-27632 (5 µM; Wako), and heparin (1 µg/ml).

Established ESC and TSC lines were adapted for feeder-free conditions. Briefly, ESCs were placed onto gelatin-coated dishes with ESC medium lacking PD0325901 and CHIR99021, and 1 μM GSK-3 inhibitor IX (Merk Millipore) was added. TSCs were placed onto dishes coated with 15 mg/mL human plasma fibronectin (Merk Millipore) (Ohinata and Tsukiyama 2014; Hirose et al. 2018). Medium was replaced every day and cells were passaged every 2 d.

For flow cytometry, cells were washed with PBS containing 1% BSA and labelled with anti-Sca-1 (D7)–PE (1:10 dilution; Miltenyi Biotec, 130-102-832). PE-positive cells were sorted by fluorescence-activated cell sorting (FACS). For differentiation assays, sorted TSCs were cultured off-feeder in CDM lacking recombinant mouse basic FGF, recombinant human activin A, XAV939, Y-27632, and heparin for 4 days prior to downstream analysis.

### Knockdown of *Setdb1* in TSCs

Oligonucleotides encoding shRNA against *Setdb1* were designed (forward: 5′ -GATCCCCGAACCTATGTTTAGTATGAACGTGTGCTGTCCGTTCATACTAAACATAGGTTCTTTTTGGAAAT-3′; reverse: 5′ -CTAGATTTCCAAAAAGAACCTATGTTTAGTATGAACGGACAGCACACGTTCATACTAAACATAGGTTCGGG-3′), annealed, and cloned downstream of the Tet operator in the pENTR4-H1tetOx1 vector. The shRNA cassette, including the Tet operator, was subsequently transferred into the lentiviral expression vector CS-RfA-ETBsd. For viral production, the resulting plasmid was transfected into 293FT cells, and viral supernatants were collected. TSCs were infected with the shRNA-expressing lentivirus in the presence of polybrene. After 24 h, the culture medium was replaced with fresh medium, and blasticidin selection (10 mg/ml) was initiated. Cells were harvested two days after the onset of selection for protein and chromatin analysis. Knockdown efficiency was assessed by Western blotting, and H3K9me3 levels at target loci were analyzed by ChIP.

### IVF and Embryo Transfer

Female mice were superovulated as described previously (PMID: 26632610). Cumulus– oocyte complexes (COCs) were collected from the oviducts 15–17 h after hCG injection and used for IVF. COCs obtained from superovulated females were placed in 80-µL droplets of human tubal fluid (HTF) medium containing 1.25 mM reduced glutathione (GSH). Spermatozoa collected from the epididymis were preincubated for 50 min in 100-µL droplets of HTF. An aliquot of the sperm suspension was then added to the HTF drops containing the COCs to initiate insemination. After incubation for 3–4 h, morphologically normal fertilized embryos were washed and cultured in CZB medium. All procedures were performed at 37 °C under 5% CO₂ in air.

Embryos that had reached the 2-cell stage after 24 h of culture were transferred into pseudo-pregnant females. Pseudo-pregnant ICR females were generated by mating with vasectomized ICR males, and the day a vaginal plug was observed was designated as embryonic day (E)0.5. Two-cell embryos were transferred into the oviducts of E0.5 pseudo-pregnant ICR females (8–12 weeks old) under anesthesia induced by intraperitoneal injection of a mixture of anesthetic agents (0.3 mg/kg medetomidine, 4.0 mg/kg midazolam, and 5.0 mg/kg butorphanol). On E19.5, fetuses were recovered by Caesarean section following cervical dislocation.

#### Histological analysis

Paraffin-embedded blocks of the placental tissue was prepared using an automatic embedding machine (Tissue-Tek VIP6, SAKURA FineTek) following the protocol below: 70% Ethanol for 1 hour, 80% Ethanol for 1 hour,90% Ethanol for 1 hour, 95% Ethanol for 1 hour, 99% Ethanol for 1 hour, 100% Ethanol for 1 hour (wice), Xylene for 1 hour (twice), Xylene for 15 minutes, Paraffin for 30 minutes (four times). I created hematoxylin and eosin (HE) stained slices of paraffin-embedded tissue using the following protocol: Xylene: 10 minutes (3 times), 100% Ethanol: 5 minutes (3 times), 95% Ethanol: 5 minutes, 75% Ethanol: 5 minutes, Rinse with tap water: 5 minutes, Rinse with distilled water: 5 minutes, Hematoxylin staining solution: 4 minutes, Rinse with tap water, Eosin staining solution: 2 minutes, 70% Ethanol, 95% Ethanol for 15 minutes,100% Ethanol for 15 minutes, Xylene for 15 minutes. I performed PAS staining according to the manufacturer’s instructions for the Periodic Acid Schiff (PAS) Stain Kit (SciTek Laboratories). The slides of the tissue sections were captured using Nanozoomer (Hamamatsu Photonics).

#### Native ChIP

Native ChIP assays were performed as described previously (Fukuda et al. 2021). Mouse monoclonal antibody against H3K9me3 (2F3) and H3.1 were used.

#### Preparation of ChIP-seq library

The ChIP DNA was fragmented by Picoruptor (Diagenode) for 10 cycles of 30 s on, 30 s off. Then, ChIP library was constructed by KAPA Hyper Prep Kit (KAPA BIOSYSTEMS) and SeqCap Adapter Kit A (Roche) according to manufacturer instructions. The concentration of the ChIP-seq library was quantified by KAPA Library quantification kit (KAPA BIOSYSTEMS). ChIP sequencing was performed on a HiSeq X platform (Illumina).

#### ChIP-seq analysis

Adaptor sequences and low quality bases in reads were trimmed using Trim Galore version 0.3.7 (http://www.bioinformatics.babraham.ac. uk/projects/trim_galore/). Then trimmed reads were aligned to the mouse GRCm38 genome assembly using bowtie version 0.12.7 (Langmead et al. 2009) with default parameters. Duplicated reads were removed using samtools version 0.1.18 (Li et al. 2009). Differential peaks were identified using *diffBind* (P-value<0.01) (Ross-Innes et al. 2012) based on narrow peaks called by MACS2 (Zhang et al. 2008). We used *Hiddendomains* (Starmer and Magnuson 2016) to identify H3K9me3 domains. Each 2 kb region was annotated as enriched or depleted H3K9me3 and continuous H3K9me3 enriched 2 kb regions was connected as H3K9me3 domain with some modifications as previously reported (Fukuda et al. 2021): the option of Bin size and max.read.count was 2000 bp and 150, respectively. Read number in each bin was normalized by the following formula to adjust difference in read number among samples. Normalized read number = read number × 30,000,000/total read number. For the hidden Markov parameter to determine enriched and depleted states, the average parameter among chromosomes rather than the parameter calculated by each chromosome was used. We merged all H3K9me3 domains for each cell type, then divided domains that were larger than 100 kb into 100-kb segments. Subsequently, we counted the read numbers mapped to each H3K9me3 domain using FeatureCounts (Liao et al. 2014) and identified differentially enriched H3K9me3 domains using Limma (FDR < 0.1).

#### Preparation of RNA-seq library

500 ng of total RNA was used for RNA-seq library construction. RNA-seq library was constructed by KAPA mRNA Hyper Prep Kit (KAPA BIOSYSTEMS) and SeqCap Adapter Kit (Roche) according to manufacturer instructions. The concentration of the RNA-seq library was quantified by KAPA Library quantification kit (KAPA BIOSYSTEMS). mRNA sequencing was performed on a HiSeq X platform (Illumina).

#### RNA-seq analysis

Adaptor sequences and low quality bases in reads were trimmed using Trim Galore version 0.3.7 (http://www.bioinformatics.babraham.ac. uk/projects/trim_galore/). The trimmed reads were mapped to the mouse GRCm38 genome assembly using TopHat v2.1.1 with –g 1 (Trapnell et al. 2009). After read mapping, the number of reads mapped in genes or repeats was counted by TEtranscripts (v1.4.11) with default parameters (Jin et al. 2015). We used Limma (Ritchie et al. 2015) to identify differentially expressed genes and transposons, focusing on those with an average reads per million (RPM) of 0.1 or higher. Differential expression was defined as having a false discovery rate (FDR) less than 0.1. In the case of placenta samples, we included the expression level of the *Hbb* gene as a covariate to account for potential red blood cell contamination. We used The Integrative Genomics Viewer (IGV) (Thorvaldsdottir et al. 2013) to visualize NGS data and ShinyGO v0.7.7 (Ge et al. 2020) for Gene ontology enrichment analysis.

#### Western blotting

Western blot analysis was performed as described previously (Fukuda et al. 2021). Briefly, cells were suspended in RIPA buffer (50mM Tris-HCl (pH 8.0), 420mM NaCl, 0.5% sodium deoxycholate, 0.1% Sodium dodecyl sulphate, 1% NP-40) and sonicated. The extract was incubated for 30 minutes on ice, and then incubated at 95°C for 5 minutes. The extract was loaded and run on SDS-PAGE gel as standard protocols. For histone proteins, intensity was analyzed by OdysseyR CLx Imagins System(LI-COR).

#### Data access

All raw and processed sequencing data generated in this study have been submitted to the NCBI Gene Expression Omnibus (GEO; https://www.ncbi.nlm.nih.gov/geo/) under accession number GSE308515 and GSE308516.

## Acknowledgments

We thank the staff of the Support Unit for Bio-Material Analysis (BMA) at the RIKEN Center for Brain Science (CBS) Research Resources Division (RRD) for NGS library construction, DNA sequencing and flow cytometry. We also thank Keiji Mochida and Ayumi Hasegawa for conducting IVF and embryo transfer experiments. We would also like to thank our colleagues at Shinkai laboratory for their support and valuable comments. RIKEN internal research fund (Pioneering project ‘Genome building from TADs’) (to Y.S. and A.O.); Y.S. was also supported by the Japan Society for the Promotion of Science (JSPS) [for Grant-in-Aid for Scientific Research [A], JP22H00413; Grant-in-Aid for Scientific Research on Innovative Areas (Research in a proposed research area), JP18H05530]; F.K. was supported by the JSPS [for Grant-in-Aid for Early-Career Scientists, 22K15044]; A.O. and K.I. were also supported by the JSPS [for Grant in Aid for Challenging Research (Pioneering) 24K21266] and [for Grant-in-Aid for Transformative Research Areas (A) 23H04956], respectively. Funding for open access charge: Japan Society for the Promotion of Science.

## Author contributions

K.F. and Y.S. designed and conceived the study. K.F. and Y.S. supervised the study and interpreted the data. C.S. performed molecular and cellular experiments and generated the ChIP-seq and RNA-seq libraries. K.F. performed informatics analysis of generated NGS data. K. I., M. H., A. O., and S. M. established TS cells and performed in vitro fertilization as well as analyses of oocyte and placental sections. M. K. analyzed the placental phenotypes. K.F. and Y.S. wrote the manuscript and prepared figures. All authors read, discussed, and approved the manuscript.

